# Contextual Science and Genome Analysis for Air-Gapped AI Research

**DOI:** 10.1101/2025.03.21.644606

**Authors:** Gerard R. Lazo, Devadharshini Ayyappan, Parva K. Sharma, Vijay K. Tiwari

**Affiliations:** USDA ARS Western Regional Research Center, Albany, CA 94710 USA; North Carolina State University, Raleigh, NC 27606 USA; University of Maryland, College Park, MD 20742 USA

## Abstract

Provided here is a study of large language models (LLMs) and retrieval augmented generation (RAG) frameworks in air-gapped environments for genome research on small grain crops. We developed two main applications: (1) a RAG-based system for contextual analysis of scientific literature, collecting over 5,000 PDFs on wheat pathogens, and (2) a GFF3 file analysis tool called Genoma that enables exploration of genome annotation through an interactive interface. Using the open-source framework Ollama, we compared the performance of multiple LLMs including Llama3.1, Deepseek-r1, and Qwen2.5 for biological data analysis. A LightRAG approach provided semantic visualization of document relationships, while the Genoma tool offered chromosome-level insights into genome annotations. These tools demonstrate the viability of powerful AI assistance for sensitive research environments, with potential applications for pangenome analysis and gene discovery. The code provides researchers with practical solutions for implementing AI in secure settings without sacrificing analytical power.

## Introduction

Technological improvements have pushed implementation of machine learning models to the generative artificial intelligence (AI) realm that we now experience. Models provided by corporate ventures are competing to bring these resources mainstream and thus is affecting the way we do research. Provided here are some examples of how some of these resources are being applied in a biological research setting.

The GrainGenes database project (Blake *et al*., 2019; Matthews *et al*., 2003) has been in service since 1992 and continues to evolve with the changes in science and technology collecting and curating data for small grains crops such as wheat, barley, rye, and oat. These crops play an important role in food security and are under continuous development to strengthen their qualities and traits for world agriculture. The recent boom in sequencing technology has accelerated the capabilities of screening plant germplasm for crop improvement and has resulted in multiple reference genomes within the species. It is a necessity for us to have a better understanding of the genome structure and functional networks to optimally identify and select associated traits to guide improvement of grain cultivars.

Discussed here will be some dialog on how we tried to apply some of these recent AI-associated technologies to assist biological research.

## Background

The public has had access to the generative AI environment since early 2023 with the Chat-GPT 3.5 release; initially with access to a handful of extensions and plug-ins to analyze datasets. Tools have significantly advanced from the early Data-Interpreter option of Chat-GPT. In early releases the context length exchanged in prompts and responses started about 8K and have incremental expanded over time to 16K, 32K, 128K range and more. Corporate offerings have added additional trained models, with extended parameters, increased context window exchanges, and many other analytical capabilities to deal with data in many multi-modal forms. Each of the corporate ventures such as OpenAI, ANTHROP\C, Google, Meta, Microsoft, X, and the like are competing to bring these resources mainstream, with each venture addressing different factors deemed desirable to the public.

Some research is conducted in data sensitive and high security environments where strict rules and practices is invoked on how data and information is shared across commodity Internet portals. Some still question how, or whether, data and information are harvested when using corporate owned systems and how this information is ingested for future uses.

Having some models openly accessible provide the ability to benchmark and develop applications within a closed research environment, termed “air-gapped”, where programs and models can be tested without contact with the Internet. Sensitive sites, such as within the federal system, could experiment with potential applications with minimal fears of adverse impacts.

The generative AI environment provides an entry-point for the public to converse with a data resource in a familiar sense without a general need for any technical training to use the interface; however, a drawback can be that the data available may lack the context and may not provide the depth of answer specificity asked of it. Data manipulation using the Langchain architecture (Chase, H., 2022) has evolved since its introduction in bringing context to data and yet retaining the power of the generative AI environment to connect to relevant data-types and information for the users prompts in an interactive manner. These approaches have since evolved into a retrieval augmented generation (RAG) approach with various variants of its implementation.

Meta AI (Ma *et al*., 2023) provided one of the first widely used large language models (LLMs) to the public, named “LLaMa” which provided a foundation for public experimentation. Data scientists soon created API interfaces, such as Open-Interpreter (Lucas, K., 2023) to assist with data analytical tasks. Likewise, other open models multiplied, and continue to do so, in repositories such as HuggingFace.co (Wolf *et al*., 2022) to extend application innovations. Another venture, ollama.ai (Morgan, J. and Chang, M., 2023), bundled a large collection of models into a 4-bit quantized framework to provide use on CPU-as well as GPU-based computer systems. A boom in experimental uses of these technologies have made their way to the public by way of YouTube.com videos and deposition at sites like github.com.

In stark contrast to some of the model genomes used for biological research such as *Arabidopsis thaliana* (100 Mb) for dicotyledonous plants, and the rice (398 Mb) and brachypodium (280 Mb) genomes for monocotyledonous grass species, our project works with small grain crops which are known for large genome size and a propensity for polyploidy. The hexaploid wheat (*Triticum aestivum*) genome is approximately 16,000 Mb. The diploid small grain crop species range from about 5,000-8,000 Mb. The primary reference genome for wheat was the germplasm ‘Chinese Spring’ sequenced in 2018 and since then, with significant improvements of sequencing technology have bloomed in numbers in recent years. There are collections of the wheat, barley, rye, and oat genomes being deposited at database repositories including GrainGenes. Ensembl, JGI, and NCBI. However, as the germplasm of these species multiply there is a need to ascertain global relationships across the pan-genome.

A variety of software tools have been used for bioinformatic analysis of genome sequences to predict genetic attributes and features. A major starting point is to detect similarities based on the National Center for Biotechnology Information (NCBI) Genbank database using the BLAST algorithm. Nowadays, it is common practice to begin annotations of a genome using a pipeline such as MAKER2 which use a collection of tools to annotate the genome (Holt and Yandell, 2011). MAKER2 produces GFF3 gene model annotations that can be visualized in genome browsers and is compatible with nucleic acid standard formats. It has been adapted to perform well on both small and large eukaryotic genomes, from microbes to plants and vertebrates. Increasingly there is a need to port these files to visualization tools such as JBrowse (Diesh *et al*., 2023), Apollo, IGV, Savant, among others. Some of these are adapted to present pan-genome visualizations. The specifications for the GFF3 file are posted (www.ncbi.nlm.nih.gov/genbank/genomes_gff), updated and followed for reported research; however, there may be a stricter need to abide by the outlined format. Within the specifications, the first eight column must be adhered to follow sequence ontology (SO) guidelines; however, there are some liberties provided for customizing the annotations in column 9, where a multitude of attributes can be added such as other ontology terms, matches to sequence and protein databases, or other repository references and notes (figure 1).

**Figure 1.**
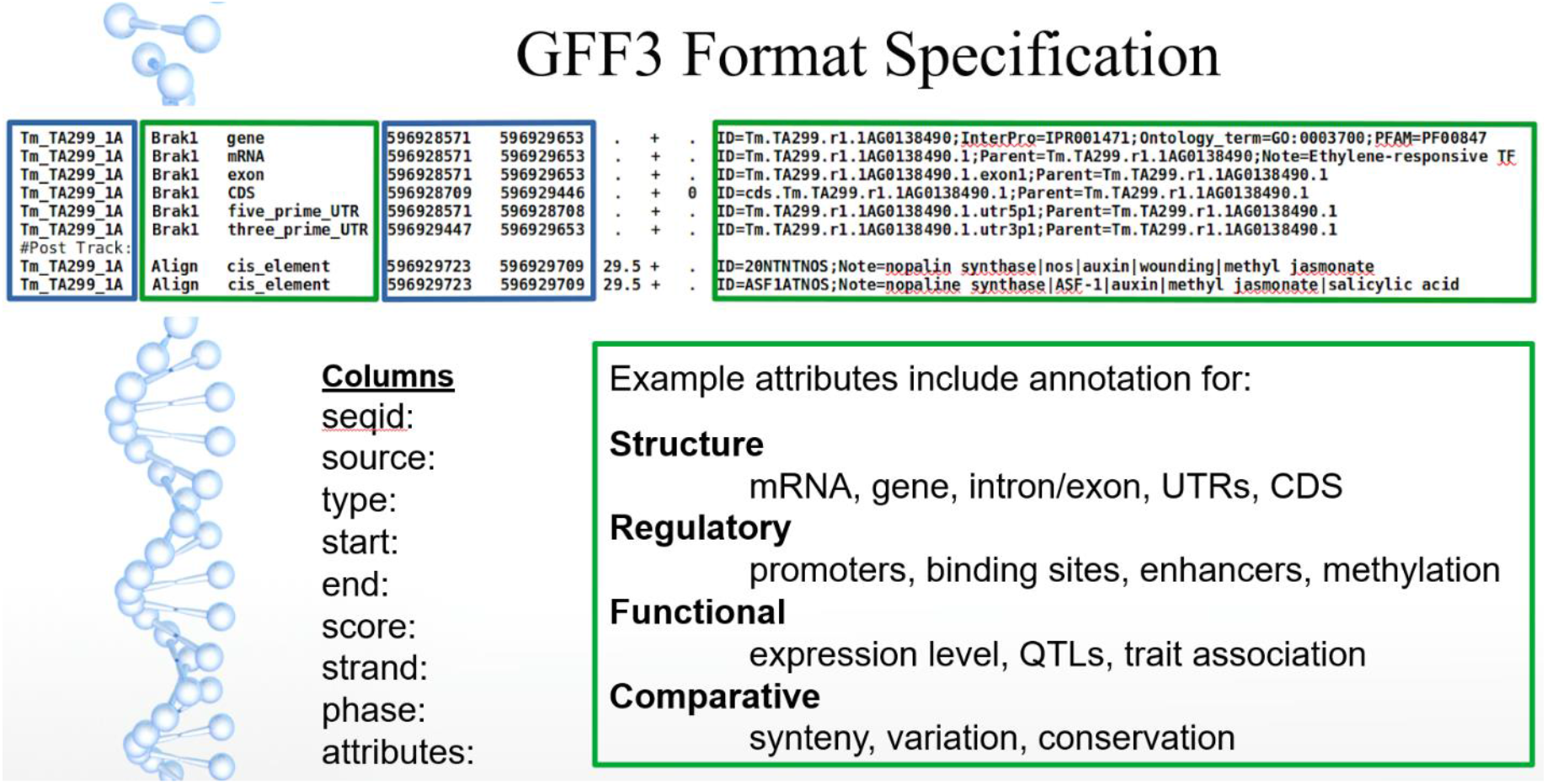
GFF3 specification. The typical series of rows covering the length of the genome region is separated in coordinated values (blue) and semantic information (green); additional qualifiers are also identified (score and condition values). Additional tracks can be created post-initial-annotation to extend information gained and generally to aid visualization, and now generative AI.

## Method

### LLMs

Hosting an LLM locally would provide an air-gapped working environment. Frameworks designed to work this way include resources from ollama.ai (for Linux, Apple, Microsoft), vLLM, LM Studio (on Microsoft). Over twenty-five fine-tuned LLMs designed for specific-purpose environments can be run with ollama (ollama.ai) with versions tuned for specific OS system memory configurations (table 2). Some models are trained at understanding computer code, others for the number of context interactions, and many based on foundation models. In the commercial realm it should be noted that OpenAI (openai.com) public models typically have a context window of 128K tokens (about 300 pages of a book) with their GPT-4 models. Early on, most of our test cases were limited to 8K tokens (including both prompts and output); however, LLMs have improved significantly, with some venturing over 1,000K. Commercial ventures often are able to provide context windows over these amounts; however, at a cost to the consumer.

### Context-aware RAG

A generative AI approach was utilized to build a knowledge-base collection for specific research topics. The design followed a Langchain framework (langchain.com) utilizing common RAG workflows (figure 2). The RAG environment provided the user an interface to converse about contextual data within a generative AI environment. To build this interface required a vector store database to index partitioned chunks of a document to be retrieved from the LLM to relate responses to a prompt.

**Figure 2.**
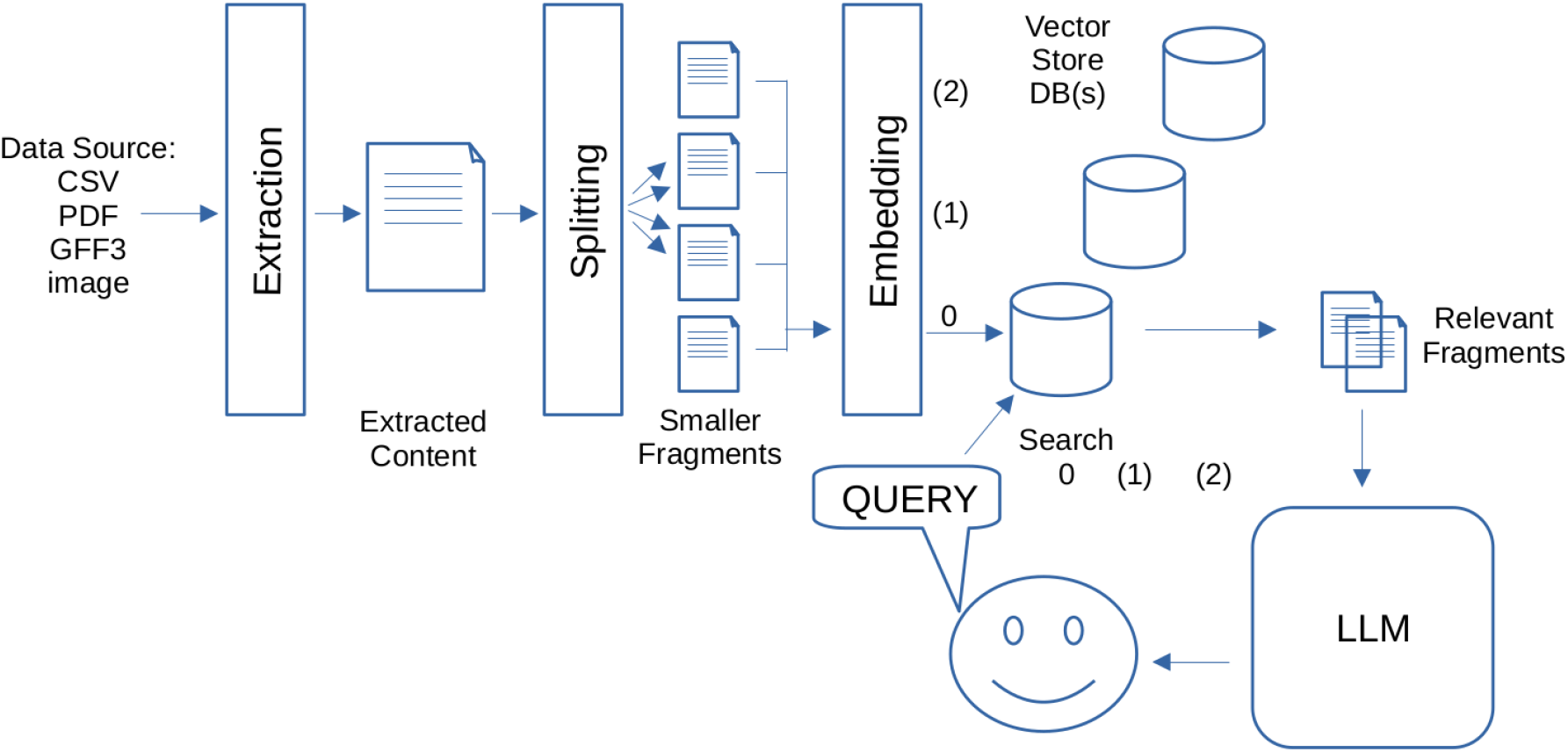
General RAG interface. Based on a langchain architecture individual data sources are extracted, split into smaller fragments, and embedded into a vector store database. In this case different vector store databases (0-2) are used for different subject topics and can be searched independently where relevant fragments pass through the LLM on to the user.

There were a number of vector store databases available; however, those available for a local system were compared. Choices included (see byby.dev/vector-databases) faiss, chroma-core, milvus-io, weaviate, vespa, vdaas/vald, qdrant, margo-ai, opensearch (fork of elasticsearch and kibana), and pgvector (extension for PostgreSQL). We primarily used chroma for these purposes.

A RAG system was established to screen PDFs for subject topics (table 1). For instance, a scientist may be interested in studying disease for their species and interactions of interest. A quick literature search can generate hundreds if not thousands of research documents that may be available over the Internet. As an example, working with wheat, there was an interest in rust, powdery mildew, bacterial blight, and blast disease.

**Table 1.**
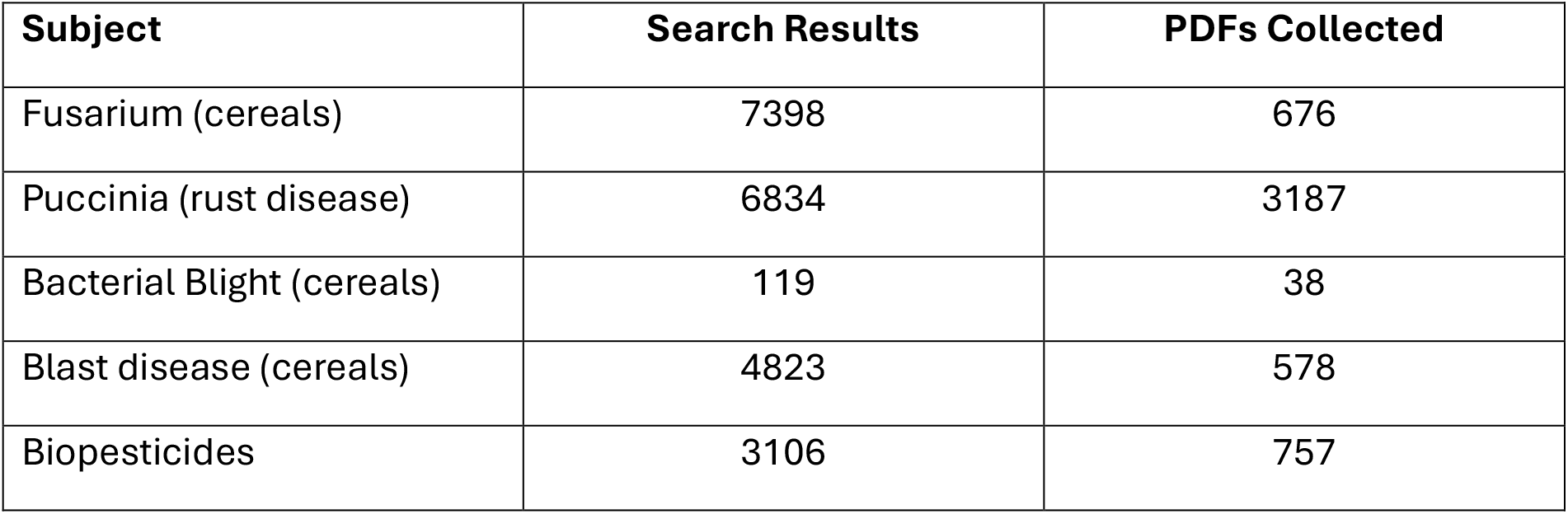
Subject topics selected for vector store database ingestion.

| **Subject** | **Search Results** | **PDFs Collected** |
| --- | --- | --- |
| Fusarium (cereals) | 7398 | 676 |
| Puccinia (rust disease) | 6834 | 3187 |
| Bacterial Blight (cereals) | 119 | 38 |
| Blast disease (cereals) | 4823 | 578 |
| Biopesticides | 3106 | 757 |

**Table 2.** Ollama LLM parameters for models tested for RAG subject topics.

| **Model** | **pa-**<br>**ram-**<br>**eters** | **context_len** | **embed-**<br>**ding_len** | **file_size** |
| --- | --- | --- | --- | --- |
| llama3:70b | 71B | 8192 | 8192 | 39G |
| phi4 (Microsoft) | 14B | 16384 | 5120 | 9G |
| mistral | 7.2B | 32768 | 4096 | 4.1G |
| qwen2.5:7b | 7.6B | 32768 | 3584 | 4.7G |
| llama3.1 | 8B | 131072 | 4096 | 4.7G |
| deepseek-r1:14b | 14.8B | 131072 | 5120 | 9.0G |
| llama3.1:405b | 410B | 131072 | 16384 | 231G |
| llama3-gradient1048K | 8B | 1048576 | 4096 | 4.7G |
| llama3-gradient:70b | 71B | 1048576 | 8192 | 39G |

For example, a collection of scientific research articles associated with the genus “Puccinia” were collected through a variety of methods. Public search engines such as Google Scholar, PubMed, Scopus, and the like yielded thousands of entries, and some were easily obtainable. Using simple web harvesting techniques PDFs were quickly accumulated, mainly by focusing on the DOI identifiers associated with the references. Over 3000 PDFs were collected for initial testing.

The data for other subject topics were also collected as PDF files and partitioned into respective directories. Each topic was ingested independently into a vector store database which was interfaced with a selected LLM to generate responses to prompts to delve into information related to the topic.

As PDFs were processed it was noted that some ill-formatted files would cause issues in the pipeline; thus, a pre-check of files using a script (pdfinfo.sh) was used to ensure file integrity. Other issues associated with the use of vector store databases also had some issues, and appropriate workarounds were implemented.

Prompting interfaces were developed using either command line or local web-browser interfaced mechanisms. In most cases the response from the prompts were programmed to relate to the top twenty responses programmatically to the LLM of choice.

### Knowledge graphs

Visualization of ingested data was available through resources such as neo4j and other methods. To adapt to a more air-gapped environment tools such as networkx were used and a LightRAG (Guo *et al*.) approach was utilized to provide general visualizations to get a sense for the quality of the data prompts and responses by noting context nodes and edges. Some GraphRAG methods consumed compute resources for large data sets and the LightRAG method provide an easy way to preview the relationships based on queried prompts.

### GFF3 prompting

Data analytics of GFF3 files were tested with early plug-ins associated with the OpenAI Chat-GPT environment (Code Interpreter), and it was presumed that it might also be applicable for the genome annotation file structure, which was primarily used for visualization of genome architecture in web-browsers. A GPT tool was even deposited at OpenAI to assist test prompting (Lazo, G.R., 2023). Early prototyping involved the use of API calls to the OpenAI GPT-3.5 model which involved using a Python Panda data-frame to read a GFF3 file and convert it to a CSV file format. Though the GFF3 was in a standard 9-column format, the ninth column at times would bear multiple annotation types which needed to be parsed out into individual columns. When interfaced with the generative AI model genome statistics and prompt associations were readily obtained. The analytical tools available from the OpenAI website directly was more illustrative than the API calls as all the data analytics were available through the commercial website. This provided a preview of what might be possible in an air-gapped environment.

## Results

RAG-based environments were developed for the subject matter PDFs and query prompts were tested against the context content using a variety of the LLMs available. As has been realized with the recent popularity of generative AI use there appears to be an art to formulating a prompt into a form which will be best ingested to provide the most appropriate response. A selection of prompts used against the “Puccinia” collection is included (table 3+). It was also noted that the models trained on more parameters usually provided the more thorough responses. Likewise, if a prompt that was asked outside of the context loaded into vector stores, the LLMs depending on their training were able to fill gaps of information based on the foundational model parameters used. New models are frequently released which purport to be more finely tuned to match larger models in a smaller footprint.

**Table 3.**
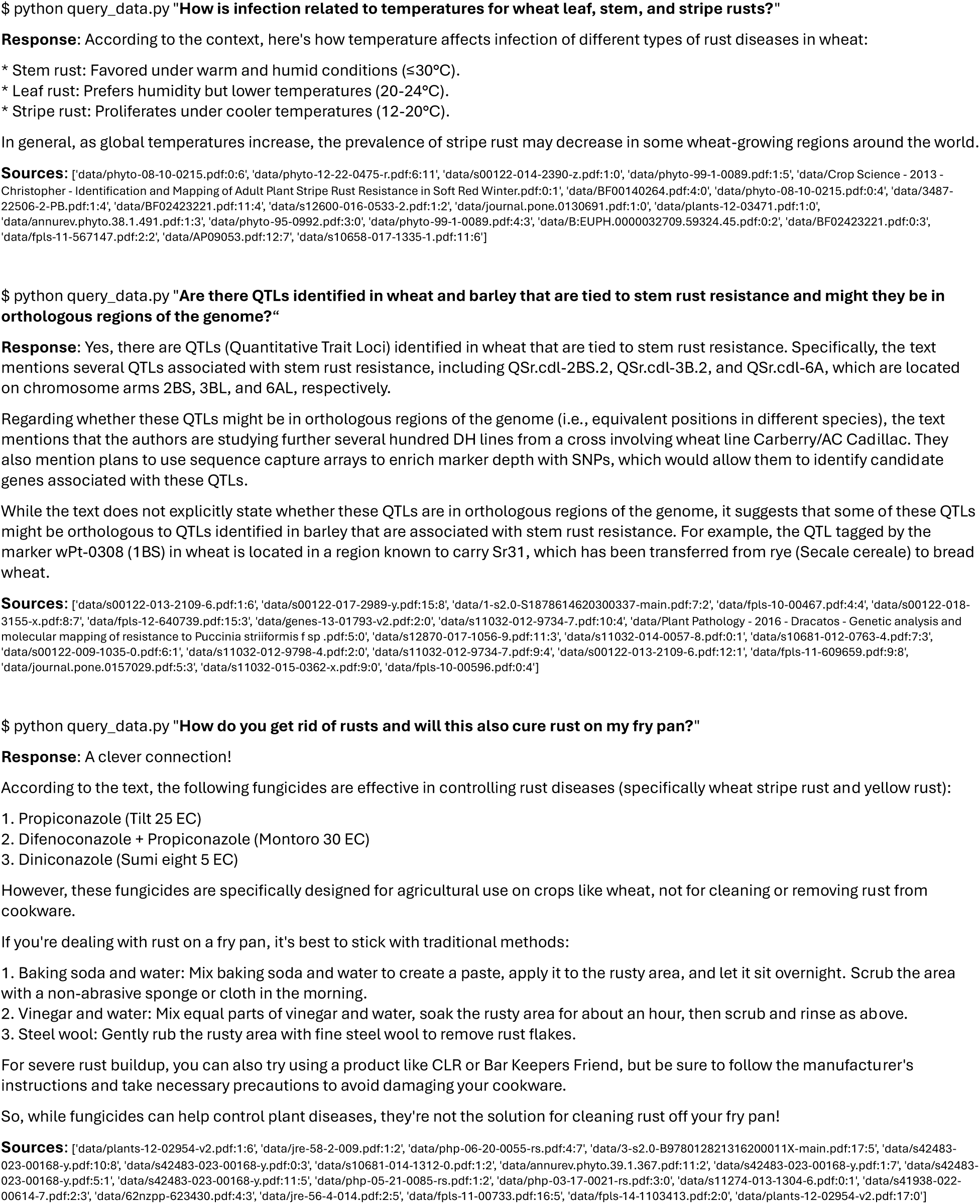
Sample prompt on command line.

Though the models run on the ollama framework did work on CPU-based computers, they were much more efficient on GPU-based machines. The conditions that you use will mainly be based on the memory that you need for the model and speed that you need to see the responses. The llama3.1 (8b) and deepseek-r1 (14b) models performed adequately on a 16GbRAM/RTX4090 GPU-based computer; however, larger, better-trained models are available for better equipped environments.

The AIRA python script for used provided a list of the top twenty documents matched in the vector store database in response to the prompt. This was factored in so that some form of validation would be confirmed upon interpreting the response. A web-based form of the prompt was also possible using a streamlit application which allowed selecting and visualizing the PDF document directly within the interface (figure 3). A quick keyword search in the document would assist confirming the validity of the top twenty pool of matches.

**Figure 3.**
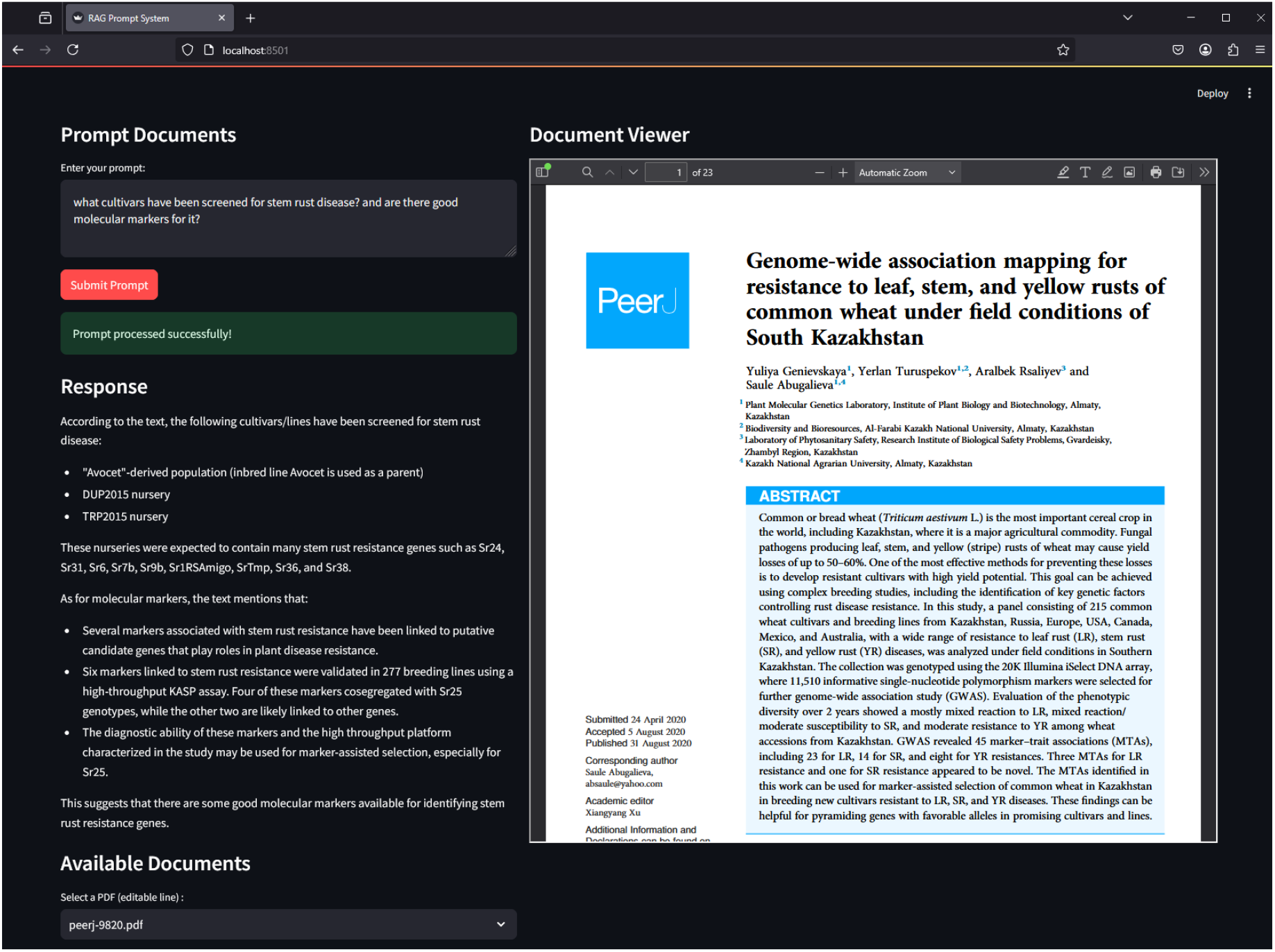
AIRA streamlit interface.

Using the LightRAG knowledge graph approach provided a simplistic overview of the complexity of terms matched with the vector databases. In a match of the top twenty matches, comparisons were visualized between the top two (#1 and #2) and bottom two (#19 and #20) knowledge graphs (figure 4). Based on the prompt the semantic features were viewable. The software did allow for some parameter alterations regarding context matches (k value) and temperature (hallucination tendency) against the vector store database which also had parameters for embedding models, context chunk size, and chunk overlap. The settings used provided satisfactory results but might be re-evaluated depending on scaling of contextual content.

**Figure 4.**
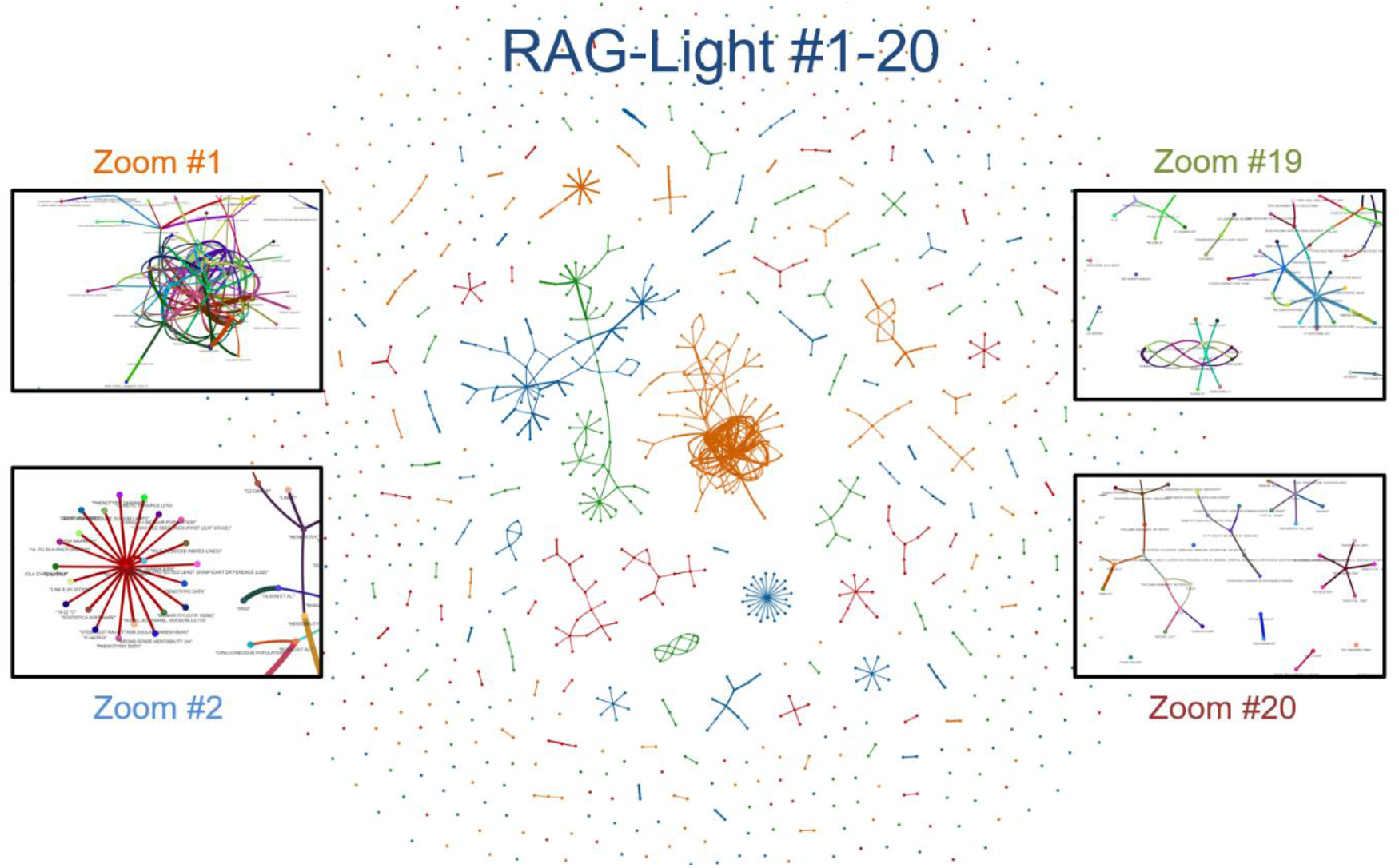
LightRAG Summary. A pool of the LightRAG treatment of the #1, #2, #19, and #20 vector store selections are shown in the middle (pattern colors in middle represent respective grouping in zoomed boxes). The top two grouping appear to have more contextual matches than the bottom two.

Early analyses on our grain genomes were limited to basically one chromosome at a time, but as context windows increase, so will the volume of data to be studied. Another work-around would be to develop agentic approaches towards working with one genome at a time. For smaller genomes, such as a microbial or model genome, perhaps a much more thorough analysis is possible (not presented here).

Most genome browsers have visualization available for gene annotations regarding transcripts, alternate transcripts, BLASTX matches, alignments with cDNAs, repetitive DNA classifications, T-DNA insertions, and the like. In a survey of over sixty available GFF3 files, the Column 9 attributes section was not uniformly represented in all files; however, each entry appeared appropriately prefixed within the semicolon separators. In the sense that we wanted to be able to query the genome we tried to use GFF3 files that were well annotated. A recent high quality sequencing effort for Einkorn wheat provides a good test for the pipeline (Ahmed *et al*., 2023). The ingest of the data used Jupyter notebooks implementing the Python Panda module to bring in data as data frames. An attribute segregation approach was implemented for Column 9 data to further place related data prefixes to be parsed under a single header. In our case, column 9 contained data for ID:, Parent:, InterPro:, Ontology_term:, PFAM:, and Note: which generally contained the semantic terms of the gene annotation.

A streamlit interface was used for creating the data frames, filtering, and prompting the content via a local LLM model. The Genoma Genome Feature Finder provides a browser interface with access to up-loadable GFF3 files which can be prompted using select LLMs. The models selected were Llama3.1, Deepseek-r1:14b, and Qwen2.5:7b; each appeared to have features which performed best depending on the prompts used. The display included tabs to different topics, each with different functionality: Genome Overview, Data Explorer, Gene Ontology, Query, Analyses, and Code Editor.

### Genome Overview

When one chromosome is selected the column 3 features are assessed and a graphical tabulation is displayed along with some minor statistics for total features, total genes, total mRNAs, average gene length, and feature density per Mb (figure 5).

**Figure 5.**
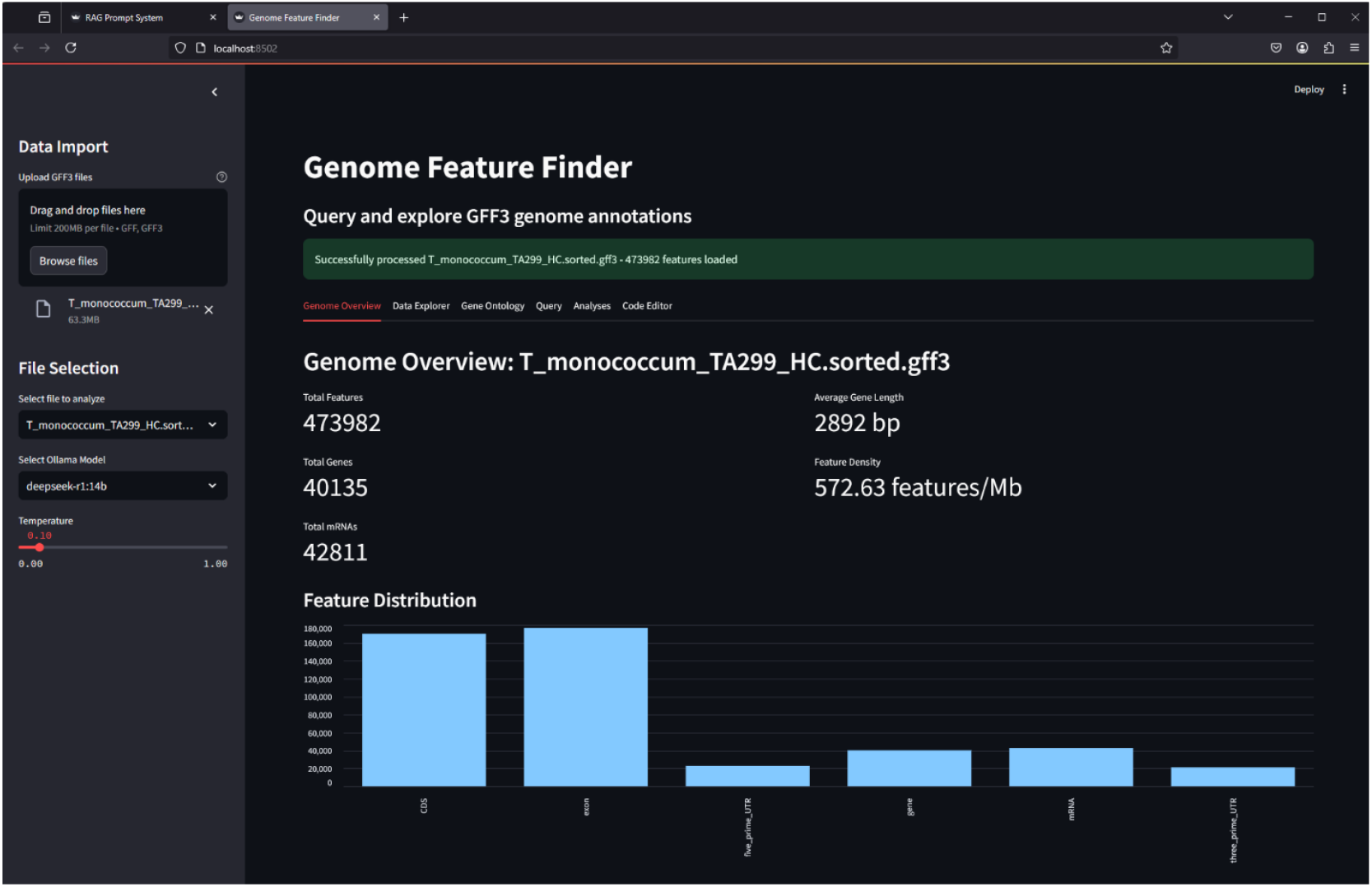
Genoma GFF file feature summary on streamlit.

### Data Explore

This tab provides a general sample data view of the GFF3 file in its raw form with a column descriptor index. An additional search and filter feature allows inspecting the file for navigable attributes of particular interest to the user. And a tab is provided to see the attributes defining the annotated gene IDs selectable from a drop-down menu.

### Gene Ontology

In many instances a user may be interested in certain attributes associated with gene ontology (GO) terms; a search can be performed using either the term ID to determine its meaning, or a semantic search term to find related GO: terms. This was an exception where an Internet connection was used to search for terms; however, creating a local database instance would obviate this need.

### Query

This area was used to directly prompt a LLM to retrieve information; depending on the prompt used different LLM were used in these instances. Some errors were sometimes thrown, but the correct answer was contained within. Software revision may correct this over time (figure 6).

**Figure 6.**
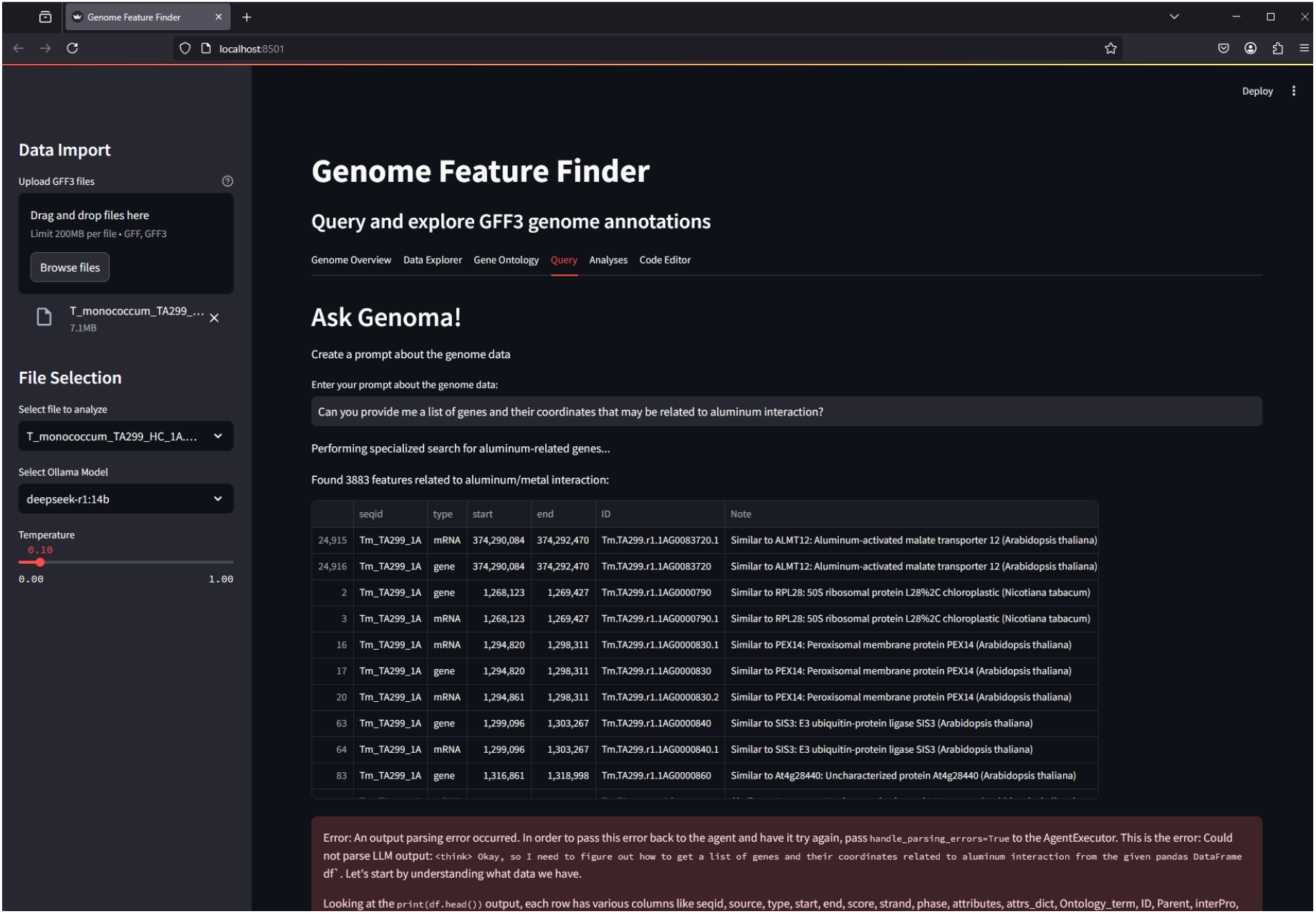
Genome GFF LLM query prompt on streamlit interface.

### Analyses

Making use of the panda data frames capabilities, some of the most common summary queries for a chromosome were placed here. The selections were selected from a drop-down menu and the analysis executed and displayed.

### Code Editor

For analyses not covered in the previous tab, a code editor window was provided for the user to attempt their own queries; a few examples are provided to preview the syntax needed (figure 7).

**Figure 7.**
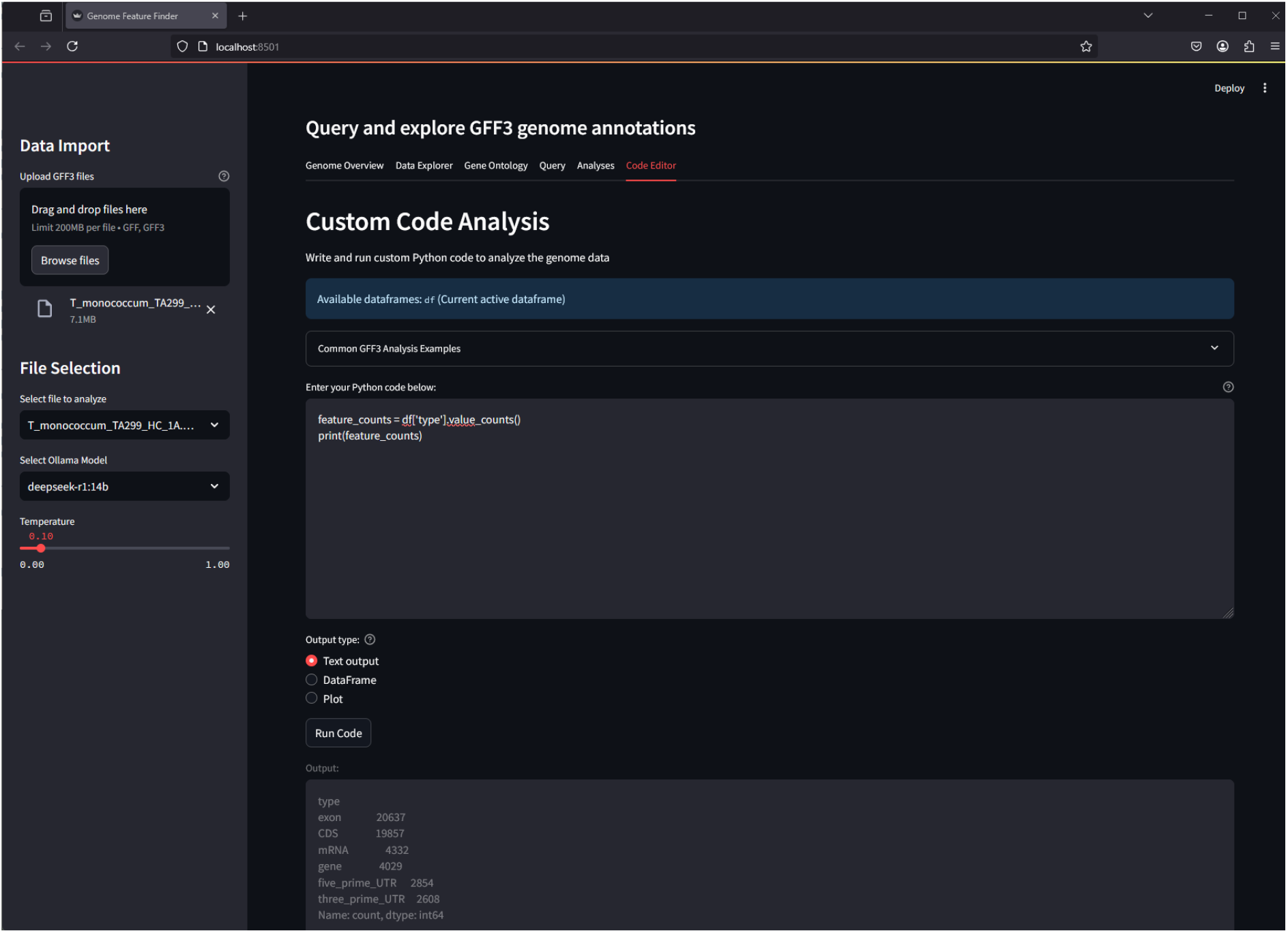
Genoma GFF custom dataframe code analysis.

## Discussion

We have provided a couple of environments to survey collections of literature topics and also a tool to examine genome annotation files. In these efforts we have utilized LLMs in an air-gapped setting utilizing generative AI prompts within an ollama.ai framework.

The literature resources can be used as a platform to have a discussion with your saved literature, basically serving as your bookshelf or file cabinet. It would be good source for brainstorming based on what you as an informed scientist should know, and at times the vector stores may point to something that you may have lost track of or may point to unique connections which were not seen before. A potential for discovery may be set to brew.

The main goal was to sample a variety of LLMs and determine constraints on the context lengths regarding volume of data and tabular-style data as was exported with the GFF3 file format. Having the capabilities to merge contextual literature with genome data can provide unique investigative approaches providing an opportunity to navigate traits associated with the genomes. Similar applications were tested in corporate offerings of OpenAI, Google, and ANTHROP\C which offered in some cases context length beyond 1 million. Paid services do offer more; however, open-source offerings are improving over time.

We have made use of many of the common tools made available as public resources for use of LLM to prompt vector store databases and data frames. The use of these tools may have some flexibility in the forms that they are used. A brief description of the methods used for collection of literature might be prerequisite to provide meta-data to the scientist to be more familiar with the terms or keywords used to build informational searches, so that RAG resources can be used and expanded upon as needed.

The use of these tools in an air-gapped system may provide a modicum of security for the user if needed; however, may be adaptable for open network connected informational environments. It was noted that there were some limitations based on the computer system specifications used. The performance was basically regulated based on the use or non-use of GPUs and limited somewhat by the memory available on the machine used. When vendor supplied services were available through Internet connections the limitations were not as dramatic as vendors were able make available additional analytical tools. So basically, one would need to realize the limitations of their work environments.

### Future

Perhaps vector store databases themselves might be shared as context-aware prompting interfaces and independently have a matching mechanism to the search engine linkable resources to protect journal offerings. Users would retrieve PDFs independently. Ultimately open literature archives may serve as new searchable resources, similar to the way BLAST databases are used for nucleic acid/protein alignments.

Though the corpus of data was extensive in some cases, there may be some limitations when working within a closed personal system. In general, the resources used were to be applied to one’s own personal collections, but the limits can still be pressed and the use of visualization through knowledge graphs may help keep a wariness of the complexity of the data. Though not exploited within these contexts, another use of knowledge graphs could be applied to better resolve relationships between the data. Limitations may rely on the capabilities of the users’ machine based on use of a GPU or not and the memory on the system.

Langchain agents and other such workflows are in development (*i*.*e*. LangGraph) to interface and to re-task prompts into other jobs, such as other vector stores or tools to create tailored agent informational responses. An agent would be able to discern the context of queries and relate to different primary keys of the GrainGenes database, for our instance, and relate contextual datatypes for genes, traits, germplasm, and the like, until the query string is complete, or another prompt is initiated. At some future point it is hopeful to relate pathogen race typing to germplasm host differentials to determine if a knowledge graph can relate germplasm and genomes to potential gene-for-gene interactions.

### Final Thoughts

We have provided a couple tools for the researcher which may aid in organizing personal data collections in the form of PDF files resources linked to LLMs in an air-gapped environment. For the interest of genome organization an additional tool has been presented to make use of genome annotation files in GFF3 format. Within this interface limited use of the LLM connections was shown possible, yet additional features allowed perusing GFF3 file content apart from their intended use in web-browser visualization.

It was noticed that there was some disparity in the standardization of the column 9 details of the GFF3 file format. Perhaps the use of the tools will help ascertain more conventional patterns to follow to create a more flexible environment to decipher genome annotations. As agents and context windows grow so will the ability to create pangenome analysis workflows.

While the air-gapped applications may not quite match what is available from corporate sponsors, it is getting there. As more and more tools are made available to the public, we can only hope to advance the technologies for the average user. As we propose and develop tools to assist our research, the AI tools will continue to aid us by improving, even coding, the software we develop, and the interfaces will continue to add different tools to agentive environments for which we may be able to conduct full investigations with a simple prompt.

## Data Availability

Computer code associated with this study will be made available or linked from resources at: github.com/USDA-REE-ARS (currently at /grlazo/AIRA, /grlazo/AIRA_Genoma, and /grlazo/AIRA_LightRAG).

## Acknowledgements

This work was supported by USDA ARS CRIS# 2030-21000-056-00D. **DA** was sponsored as a 2023 AI-COI/SCINET Graduate Research Internship. **PKS** was sponsored as a postdoctoral scientist under a cooperative agreement between **GRL** at USDA and **VKT** at University of Maryland; NACA#58-2030-1-029.

## Notes

### Competing Interest Statement

The authors have declared no competing interest.

